# Mcc1229, an Stx2a-amplifying microcin, is produced *in vivo* and requires CirA for activity

**DOI:** 10.1101/2021.07.19.453020

**Authors:** Erin M. Nawrocki, Laura E. Hutchins, Kathryn A. Eaton, Edward G. Dudley

**Affiliations:** Department of Food Science, The Pennsylvania State University, University Park, PA, USA; Department of Microbiology and Immunology, University of Michigan, Ann Arbor, MI, USA; *E. coli* Reference Center, The Pennsylvania State University, University Park, PA, USA

## Abstract

Enterohemorrhagic *E. coli* (EHEC) strains, including the foodborne pathogen *E. coli* O157:H7, are responsible for thousands of hospitalizations each year. Various environmental triggers can modulate pathogenicity in EHEC by inducing expression of Shiga toxin (Stx), which is encoded on a lambdoid prophage and transcribed together with phage late genes. Cell-free supernatants of the sequence type (ST) 73 *E. coli* strain 0.1229 are potent inducers of Stx2a production in EHEC, suggesting that 0.1229 secretes a factor that activates the SOS response and leads to phage lysis. We previously demonstrated that this factor, designated microcin (Mcc) 1229, was proteinaceous and plasmid-encoded. To further characterize Mcc1229 and support its classification as a microcin, we investigated its regulation, determined its receptor, and identified loci providing immunity. Production of Mcc1229 was increased upon iron limitation, as determined by ELISA, *lacZ* fusions, and qRT-PCR. Spontaneous Mcc1229-resistant mutants and targeted gene deletion revealed that CirA was the Mcc1229 receptor. TonB, which interacts with CirA in the periplasm, was also essential for Mcc1229 import. Subcloning of the Mcc1229 plasmid indicated that Mcc activity was neutralized by two ORFs, each predicted to encode a domain of unknown function (DUF)-containing protein. In a germfree mouse model of infection, colonization with 0.1229 suppressed subsequent colonization of EHEC. Although Mcc1229 was produced *in vivo*, it was dispensable for colonization suppression. The regulation, import, and immunity determinants identified here are consistent with features of other Mccs, suggesting that Mcc1229 be included in this class of small molecules.

## Introduction

Enterohemorrhagic *Escherichia coli* (EHEC) are foodborne pathogens that can cause severe clinical complications, including hemorrhagic colitis (HC) and hemolytic uremic syndrome (HUS), through the production of Shiga toxin (Stx) and other virulence factors (1–3). Stx is encoded on a temperate lambdoid bacteriophage and is therefore induced via the bacterial SOS response (4–6). Certain antibiotics and DNA-damaging agents are known to trigger phage induction and increase the expression of Stx *in vivo* and *in vitro* (7, 8). In the intestinal environment, members of the microbiome and their metabolites can modulate the pathogenicity of EHEC strains in multiple ways (9). Commensal bacteria can reduce the growth and colonization of EHEC, broadly limiting virulence factor expression (10). Alternatively, strains that are sensitive to the *stx*-converting phage can be infected and thus amplify Stx production (11–13). Finally, small molecules such as bacteriocins that target EHEC can both inhibit growth and promote Stx expression by induction of the phage lytic cycle (14, 15).

Bacteriocin activity was first described nearly a century ago (16) and is widespread in *E. coli*, with up to 60% of strains identified as colicin producers in some surveys (17–19). Microcins, which have a lower molecular weight than colicins (20), are found less frequently and are not as well characterized (21). They are generally smaller than 10 kDa in size, are not SOS-induced, and are secreted by intact cells (22, 23). Foundational studies on microcin B17 (MccB17), MccJ25, and others revealed that microcins are typically expressed in stationary phase, when cells are starved for nutrients (24–26). In particular, iron-limiting conditions often stimulate microcin production (27–29). Some microcins are post-translationally modified with the addition of siderophores (30–32), and many colicins and microcins exploit siderophore receptors for entry into target cells (33, 34). Expression of bacteriocins in nutrient-poor environments can also confer a fitness advantage to producing strains, allowing them to kill their competitors and better colonize a given niche (35–37). In mouse models, for example, iron limitation can be advantageous for either pathogens (38) or probiotic bacteria (39) that produce bacteriocins.

Prior studies of the human *E. coli* isolate 0.1229 revealed that cell-free supernatants from this strain were sufficient to induce the SOS response and increase the Stx expression of EHEC (15). Microcin B17, which is encoded on a 96.3 kb plasmid in 0.1229, contributed to but was not fully responsible for SOS induction or Stx amplification (15). An additional factor with Stx-amplifying activity was localized to p0.1229_3, a 12.9 kb plasmid in the strain (15). This activity was dependent on TolC for efflux from 0.1229 and TonB for import into the target cell (15). The SOS-inducing, Stx-amplifying agent of p0.1229_3 is presumed to be a new microcin, first described in strain 0.1229 and thus designated Mcc1229. Although the chemical identity of Mcc1229 is not known, it is encoded within a 5.2 kb region of p0.1229_3 whose annotations include hypothetical proteins, an ABC transporter, a cupin superfamily protein, and domain of unknown function (DUF)-containing proteins (15).

Only a small number of microcins have been purified, and their functions in complex environments like the gut microbiome are not well defined (21). Some have theorized that the microcins prevalent in phylogroup B2 *E. coli* enhance their ability to dominate the rectal niche and colonize the urinary tract (40). 0.1229 is a phylogroup B2 isolate of sequence type (ST) 73. Other members of ST73 are notable urinary pathogens (e.g. CFT073), and the lineage carries many virulence factors that can promote colonization and persistence *in vivo* (41, 42). In 0.1229, MccB17 and Mcc1229 may serve this purpose, as they are lethal to competing *E. coli* strains (15). To elucidate the role of the putative microcin Mcc1229, we have here clarified its export, import, immunity, and regulation. We have also probed the effect of 0.1229 and its microcins in a germfree mouse model of EHEC infection.

## Results

### Stx2a levels are increased upon growth in supernatants of E. coli 0.1229 ΔB17::FRT

*E. coli* strain 0.1229 produces two microcins: microcin B17 and the lesser characterized Mcc1229 (15). To isolate the impact of Mcc1229, we deleted the microcin B17 operon by one-step recombination, generating 0.1229 ΔB17::FRT. Inactivation of both microcins was accomplished by one-step recombination of 0.1229 ΔB17::FRT with the Δ6::*cat* PCR product, which was previously designed to remove a hypothetical protein and an ABC transporter from the Mcc1229 cluster on p0.1229_3 (15). These ORFs were formerly described as Hp1 and ABC based on their predicted protein products (15). In accordance with microcin nomenclature and in reference to their role in amplifying Shiga toxin, we have assigned genes in the Mcc1229 region names that begin with *mctA* (<u>m</u>icro<u>c</u>in involved in toxin <u>a</u>mplification). Hp1 and ABC have been renamed *mctA* and *mctB*, respectively, and therefore the Δ6::*cat* deletion will be now be referenced as Δ*mctAB*::*cat*.

To determine the effect of culture conditions and to optimize production of Mcc1229 for future studies, *E. coli* 0.1229 ΔB17::FRT was grown in various liquid media. These included LB with 0%, 0.5%, and 1% NaCl (no, low, and high salt, respectively) and M9 supplemented with casamino acids, thiamine, and 0.4% of glucose, glycerol, maltose, or fructose. Stx2a amplification by Mcc1229 was determined by culturing the *stx*_2a_ *E. coli* O157:H7 strain PA2 in spent supernatants. Supernatants from LB cultures of 0.1229 ΔB17::FRT amplified Stx2a to the greatest extent (Figure 1). Amplification was dependent on Mcc1229 production, as the supernatants of 0.1229 ΔB17::FRT Δ*mctAB*::*cat* (which produced neither MccB17 nor Mcc1229) were statistically equivalent to broth controls (p = 0.9789) and induced minimal levels of Stx2a (Figure 1).

**Figure 1.**
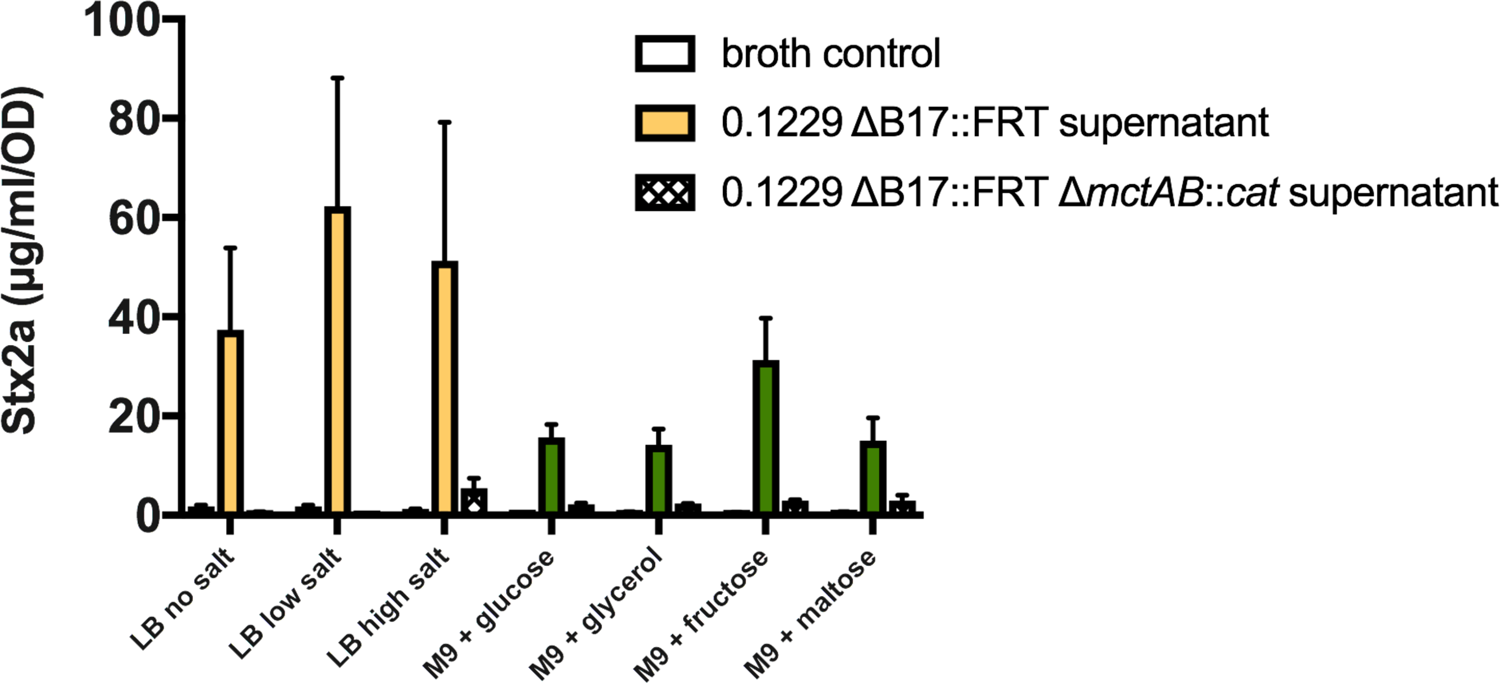
Stx2a levels are amplified by culture supernatants from rich media. Cell-free supernatants of *E. coli* 0.1229 ΔB17::FRT grown in various media were used to culture the *E. coli* O157:H7 strain PA2. Stx2a levels were measured by R-ELISA and normalized to the OD_620_ of each culture; the mean + SEM are reported (minimum n = 3). M9 medium was supplemented with 0.1% casamino acids, 0.005% thiamine, and 0.4% of the given carbon source. Cultures grown in spent supernatants of 0.1229 ΔB17::FRT are indicated by filled bars (yellow for LB, green for M9). Cultures grown in spent supernatants of the double microcin mutant, 0.1229 ΔB17::FRT Δ*mctAB*::*cat*, are given as crosshatched bars. For comparison, PA2 was grown in fresh media of the same composition, as indicated by empty bars. All such “broth control” cultures yielded less than 5 µg/ml/OD Stx2a.

### Iron suppresses Stx2a-amplifying activity

To further investigate conditions influencing Mcc1229 expression, we added the metal chelating agents EDTA or 2,2’-bipyridyl (bipy) to cultures of 0.1229 ΔB17::FRT grown in LB with high salt. Supernatants from 0.1229 ΔB17::FRT cultures grown in LB + bipyridyl amplified Stx2a levels beyond those from LB alone (Figure 2). In the presence of low concentrations (10 µM) of ferric chloride, EDTA and bipyridyl supernatants still significantly amplified Stx2a above unsupplemented LB supernatant levels (Figure 2). When FeCl_3_ levels were increased to 200 µM— equimolar to EDTA or bipyridyl—the Stx2a-amplifying effect of these supernatants was suppressed (Figure 2). In other words, excess iron negated the impact of bipyridyl on Mcc1229. By contrast, the addition of 200 µM CaCl_2_, MgCl_2_, or MnCl_2_ to 0.1229 ΔB17::FRT cultures did not reduce their Stx2a-amplifying activity (data not shown), suggesting that this effect was specific to FeCl_3_.

**Figure 2.**
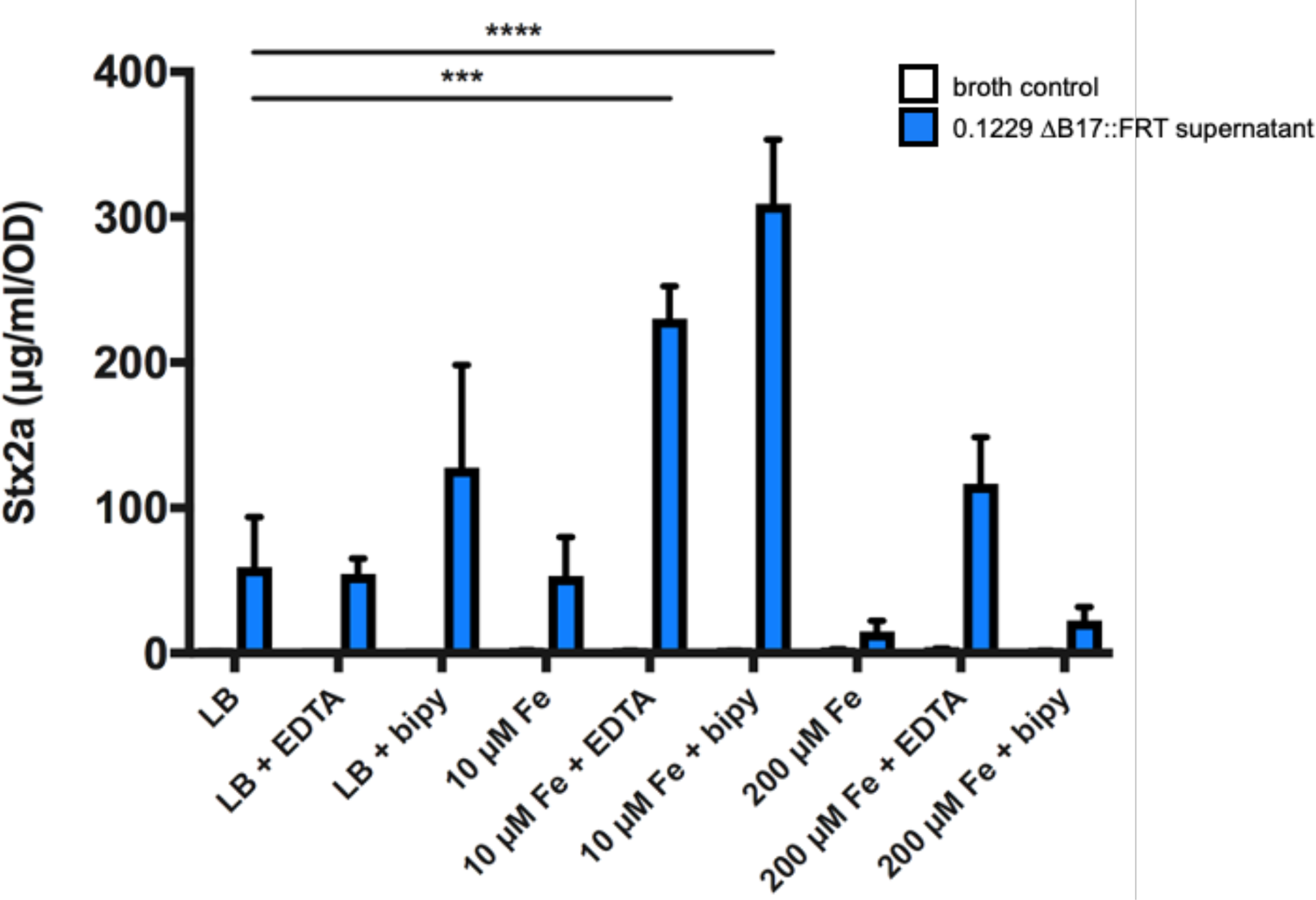
Stx2a levels are diminished when grown in supernatants from high-iron media. Cell-free supernatants of *E. coli* 0.1229 ΔB17::FRT grown in various media were used to culture the *E. coli* O157:H7 strain PA2. For comparison, PA2 was grown in fresh media of the same composition, as indicated by “broth control.” Stx2a levels were measured by R-ELISA and normalized to the optical density of each culture; the mean + SEM are reported (n = 3 in all except 10 µM Fe cultures, for which n = 2). The metal chelators EDTA and 2,2’-bipyridyl (bipy) were added to media at 0.2 mM. Statistical significance was determined by two-way ANOVA and Sidek’s multiple comparisons test, assigning LB as the standard for the broth control and 0.1229 ΔB17::FRT supernatant groups (***, p < 0.001; ****, p < 0.0001).

### mctA, mctB, and mctC ORFs are required for Stx2a amplification

Prior work demonstrated that the p0.1229_3 plasmid, and specifically a 5.2 kb region therein, was sufficient to amplify Stx2a (15). This region was moved into the medium-copy pBR322 vector, replacing the *bla* gene (15). The resulting construct is termed pBR322::mcc1229. The mcc1229 region is predicted to encode three hypothetical proteins (MctA, MctF, and MctG), an ABC transporter (MctB), a cupin domain protein (MctC), and two domain of unknown function proteins (MctD and MctE) (Figure 3a). Each of these was previously deleted by one-step recombination in the native 0.1229 strain, inserting a *cat* marker in its place on p0.1229_3. *mctA*, *mctB*, *mctC*, and *mctF* deletion mutants were significantly impaired for Stx2a amplification (15). To avoid any potential polar effects of the *cat* insertions, we here constructed markerless in-frame deletions of the same ORFs in the pBR322::mcc1229 clone instead. When the *mctA*, *mctB*, and *mctC* ORFs were deleted from this region, the resulting constructs no longer amplified Stx2a (Figure 3b). Supernatants from C600 (pBR322::mcc1229Δ*mctA*), C600 (pBR322::mcc1229Δ*mctB*), and C600 (pBR322::mcc1229Δ*mctC*) were indistinguishable from an empty vector control (Figure 3b). C600 (pBR322::mcc1229Δ*mctF*) and C600 (pBR322::mcc1229Δ*mctG*) were not diminished in their ability to amplify Stx2a (data not shown).

**Figure 3.**
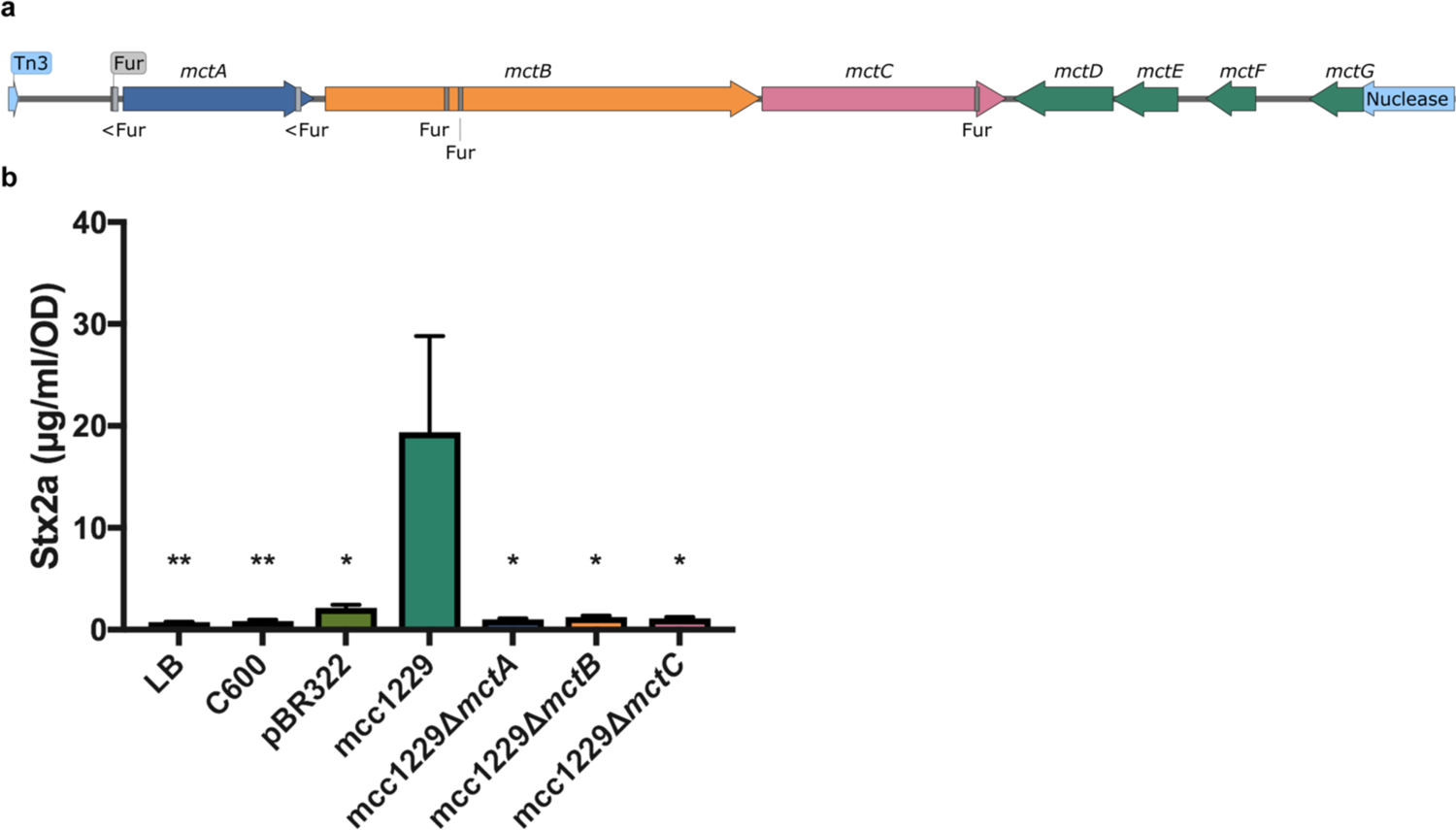
A 5.2 kb region of p0.1229_3 sufficient for Stx2a amplification encodes seven putative open reading frames and six predicted Fur binding sites. (a) Annotation of p0.1229_3 was performed by NCBI’s Prokaryotic Genomes Automatic Annotation Pipeline (PGAAP) as reported previously (15, 82). Fur sites were identified by the matrix-scan algorithm at the RSAT Prokaryotes webserver (79, 83). Those on the reverse strand are indicated by <. The map diagram was generated by SnapGene software (from GSL Biotech; available at snapgene.com). (b) The mcc1229 region of p0.1229_3 was cloned into pBR322 and was sufficient to amplify Stx2a. Supernatants from the C600 strain alone or from the empty vector pBR322 did not amplify Stx2a. Stx2a expression of PA2 exposed to filtered culture supernatants was determined by ELISA as previously described. Values that differed significantly from the mcc1229 supernatant are marked with asterisks (*, p < 0.05; **, p < 0.01). Statistical analysis was performed by one-way ANOVA with Dunnett’s multiple comparisons test.

### p0.1229_3 ORFs are in the Fur regulon

Sequence analysis of the Stx2a-amplifying region on p0.1229_3 revealed a putative Fur binding site [ataAATGATAActATTcTC, where uppercase letters indicate identity to the consensus (43)] upstream of the *mctA* open reading frame (Figure 3a). The region upstream of *mctA* was ligated into pRS551 and successfully promoted transcription of *lacZ* (Figure 4a). Promoter regions upstream of the *mctB*, *mctC*, and *mctE* ORFs were also verified in this manner (Figure 4a). Transcriptional activity of the *mctA* promoter decreased when supplemented with ferric chloride (Figure 4a), suggesting that MctA was regulated by iron. These data were supported by transcriptional analysis in 0.1229 and 0.1229 Δ*fur*::*cat*. qRT-PCR targeting *mctA*, *mctB*, *mctC*, and *mctE* regions indicated that expression of each gene was increased in the *fur* mutant (Figure 4b), consistent with a model in which the Fe-Fur complex repressed transcription of the microcin.

**Figure 4.**
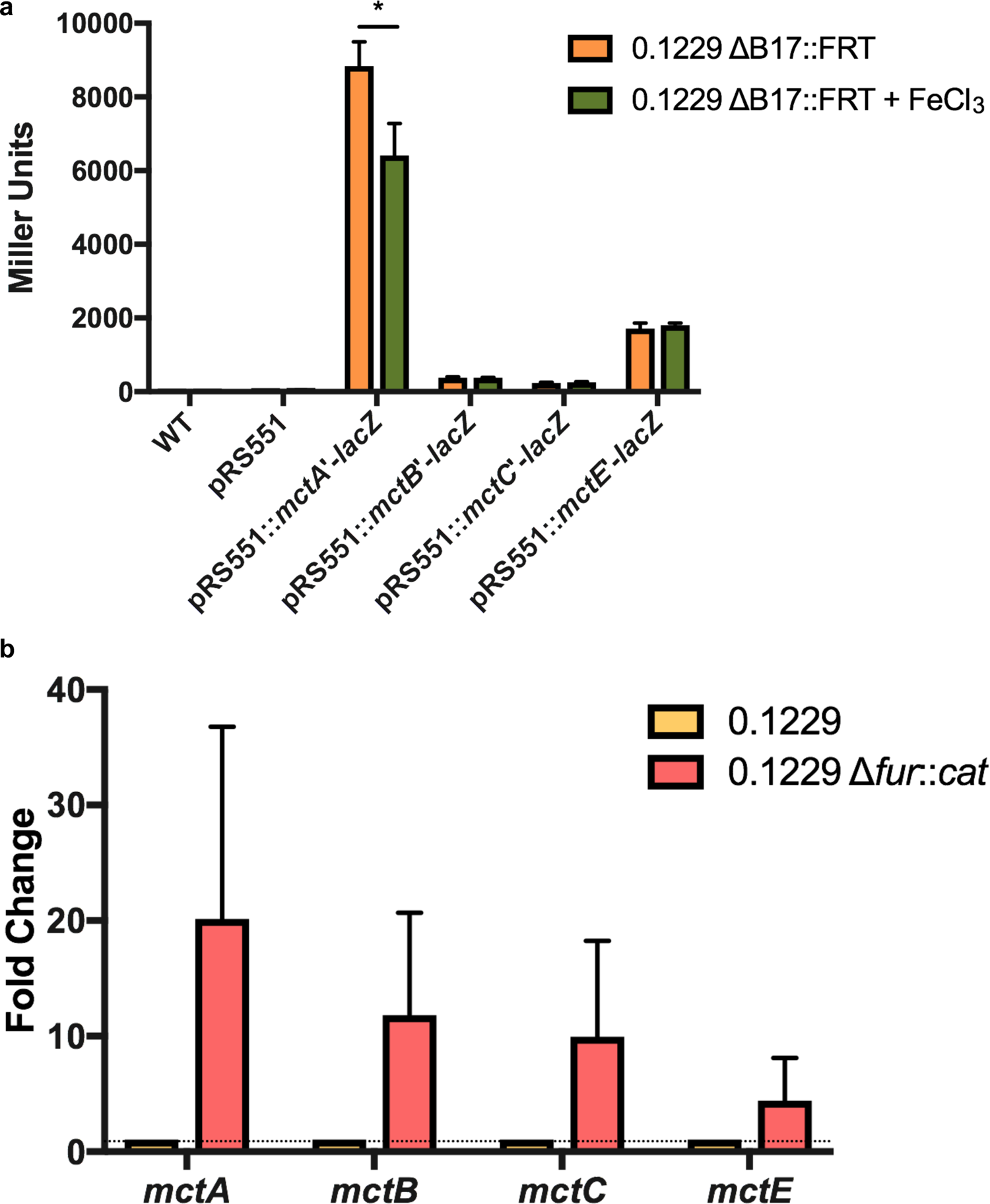
Mcc1229 transcription is iron-regulated. (a) A region upstream of the *mctA* ORF containing a putative Fur binding site was ligated into pRS551 and successfully promoted transcription of *lacZ*. Promoter regions upstream of the *mctB*, *mctC*, and *mctE* ORFs were also tested in this manner. Transcriptional activity of the *mctA* promoter decreased when the medium was supplemented with 200 µM FeCl_3_. Significance was determined by two-way ANOVA with Sidak’s multiple comparisons test. (b) RNA was extracted from 16 h LB cultures of 0.1229 and 0.1229 Δ*fur*::*cat*, converted to cDNA, and probed by qPCR for the *mctA*, *mctB*, *mctC*, and *mctE* ORFs. Gene expression relative to wildtype was determined by the ΔΔC_T_ method using the ribosomal gene *rrsH* as an internal control (84). All Mcc1229 genes were consistently upregulated in the Δ*fur*::*cat* strain but the difference was not statistically significant by two-way ANOVA.

### CirA is the outer membrane receptor for Mcc1229

To identify the receptor for Mcc1229, we investigated its entry into target cells in several ways. First, in an agar overlay assay, we showed that Mcc1229 inhibited the susceptible *E. coli* O157:H7 strain PA2, creating a zone of clearing around Mcc1229-producing colonies. Spontaneous Mcc1229-resistant mutants of PA2 that grew within the zone of inhibition were then subject to whole-genome sequencing. Three independent colonies revealed mutations that would affect CirA expression: two contained frameshift mutations in the *cirA* ORF that introduce a premature stop codon, and one carried an 86 bp deletion directly upstream of the *cirA* start codon (Table 1). These isolates (PA2.1, PA2.2, and PA2.3) were insensitive to Stx2a amplification by 0.1229 ΔB17::*cat* supernatants (Figure 5). Second, a targeted deletion of *cirA* in PA2 by one-step recombination (Δ*cirA*::*kan*) was also resistant to Mcc1229-mediated Stx2a amplification (Figure 5). Sensitivity was restored by complementation with the medium-copy number pBR322::*cirA* (Figure 5). Finally, *cirA* was confirmed as the Mcc1229 receptor using a set of indicator strains bearing mutations in known colicin receptors (44). While a wild-type indicator strain was susceptible to Mcc1229 inhibition in the agar overlay method, a *cirA* mutant was resistant to this microcin (data not shown).

**Figure 5.**
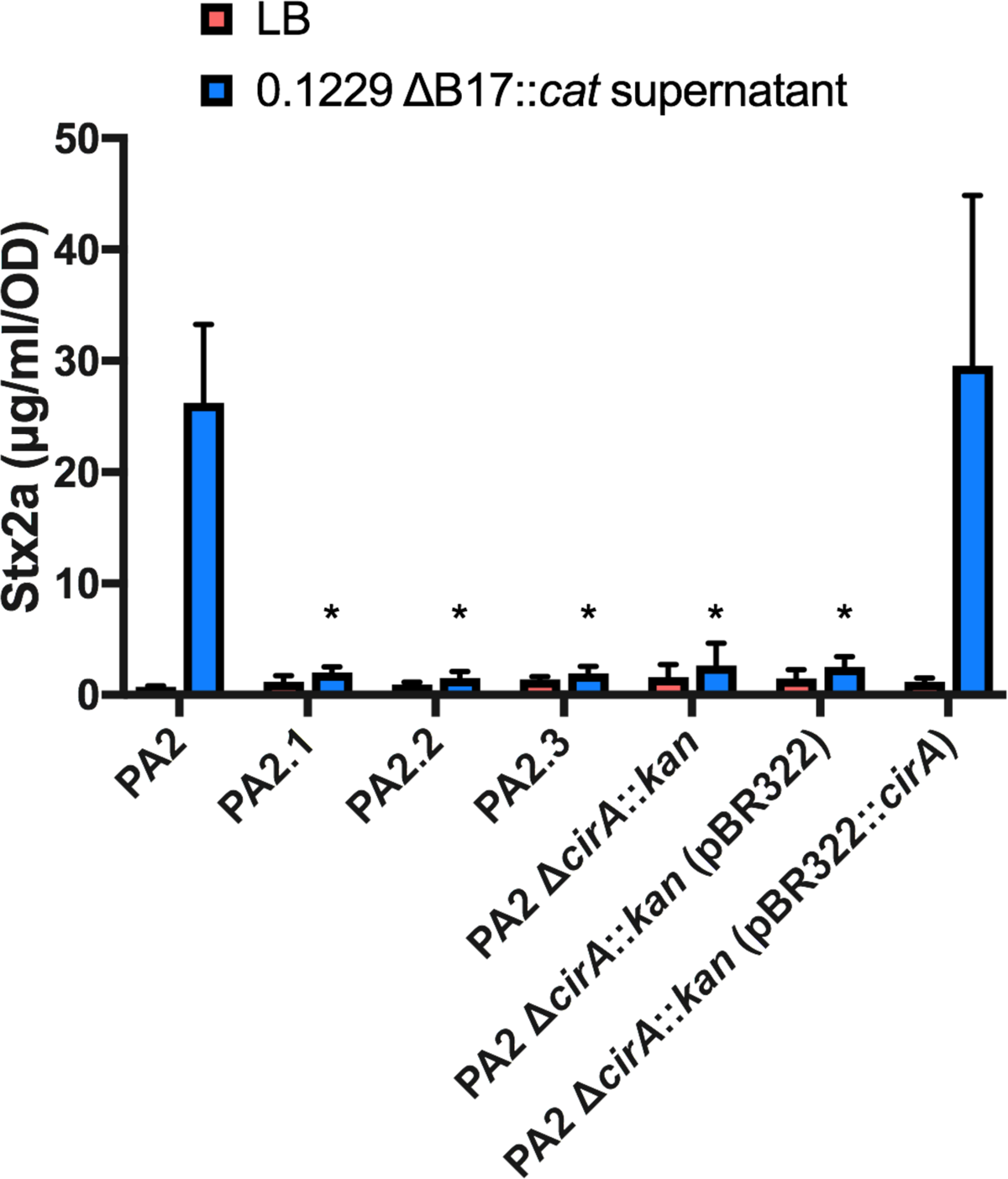
CirA is the outer membrane receptor for Mcc1229. Spontaneous mutants of PA2 were resistant to inhibition by *E. coli* 0.1229 Δ*B17*::*cat*. Colonies were isolated from within the zones of clearing and subject to whole genome sequencing to identify the source of Mcc1229 resistance. Multiple independent mutants (PA2.1, PA2.2, PA2.3) had mutations in CirA. A Δ*cirA*::*kan* mutant of PA2 is resistant to Mcc1229, and when *cirA* mutants are grown in spent supernatants of *E. coli* 0.1229 Δ*B17*::*cat*, they are insensitive to Stx amplification. Sensitivity is restored by complementation with pBR322::*cirA*. Asterisks mark significant difference between a given supernatant sample and the PA2 wildtype (two-way ANOVA, Dunnett’s multiple comparisons test).

**Table 1.**
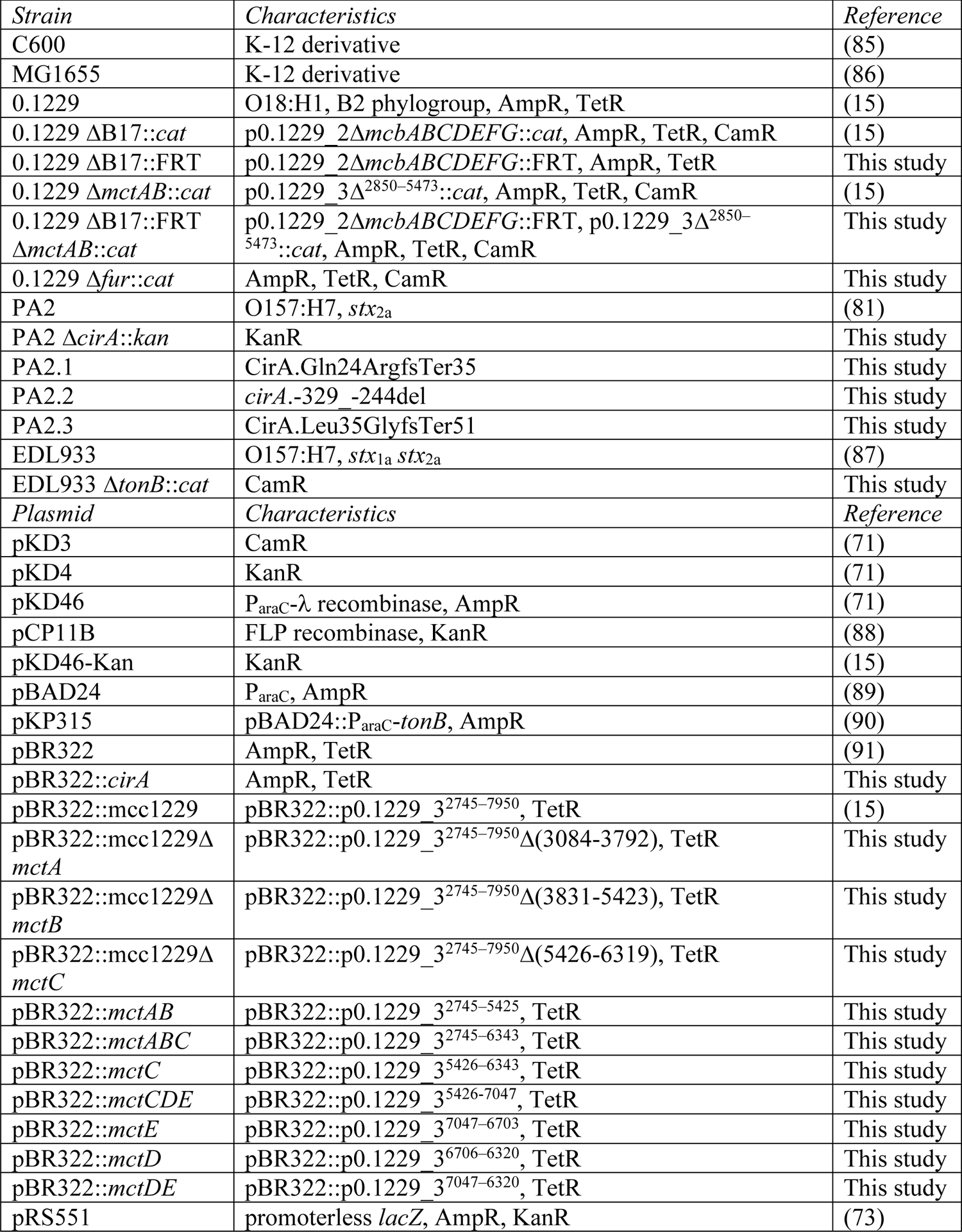
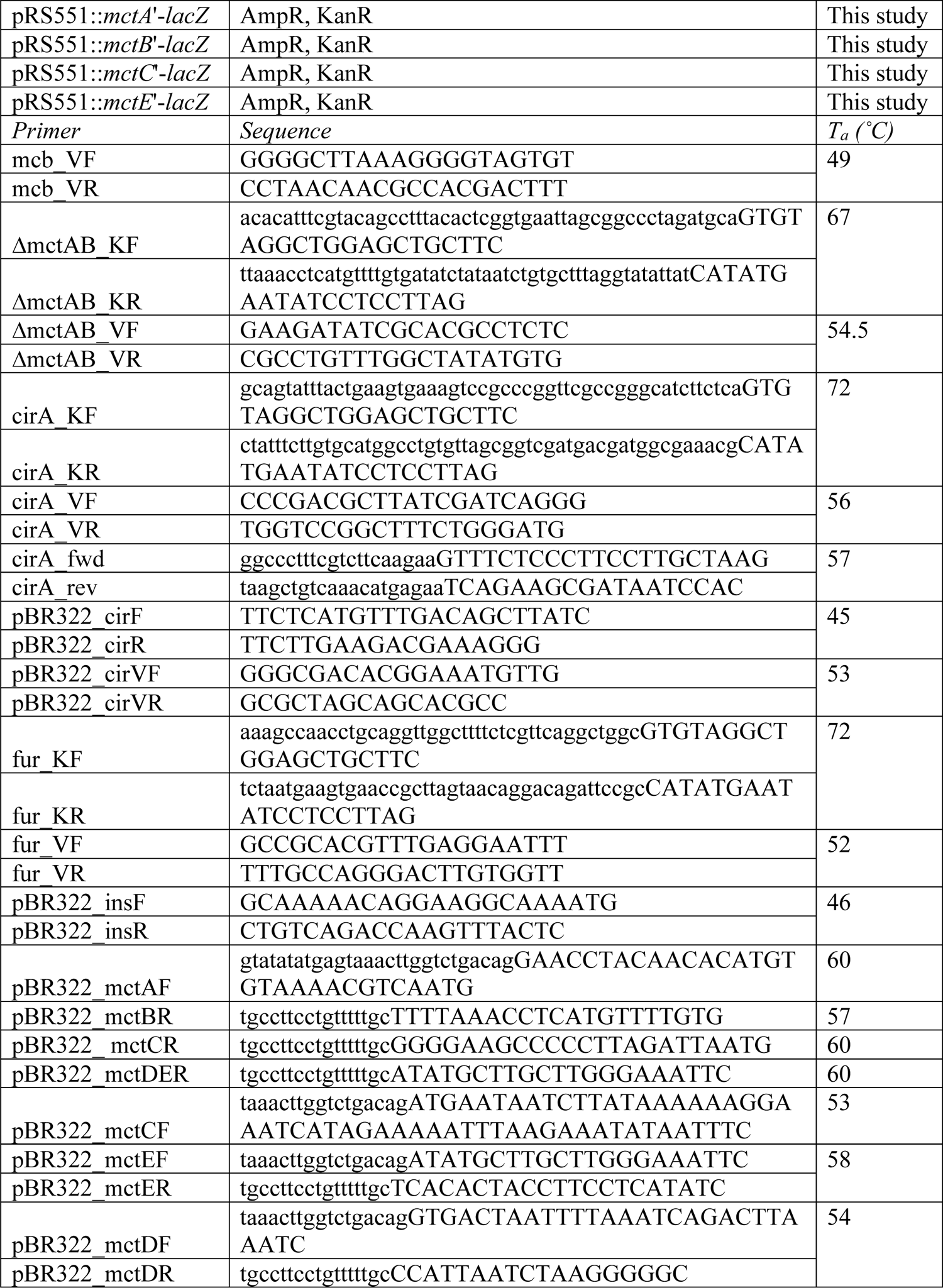
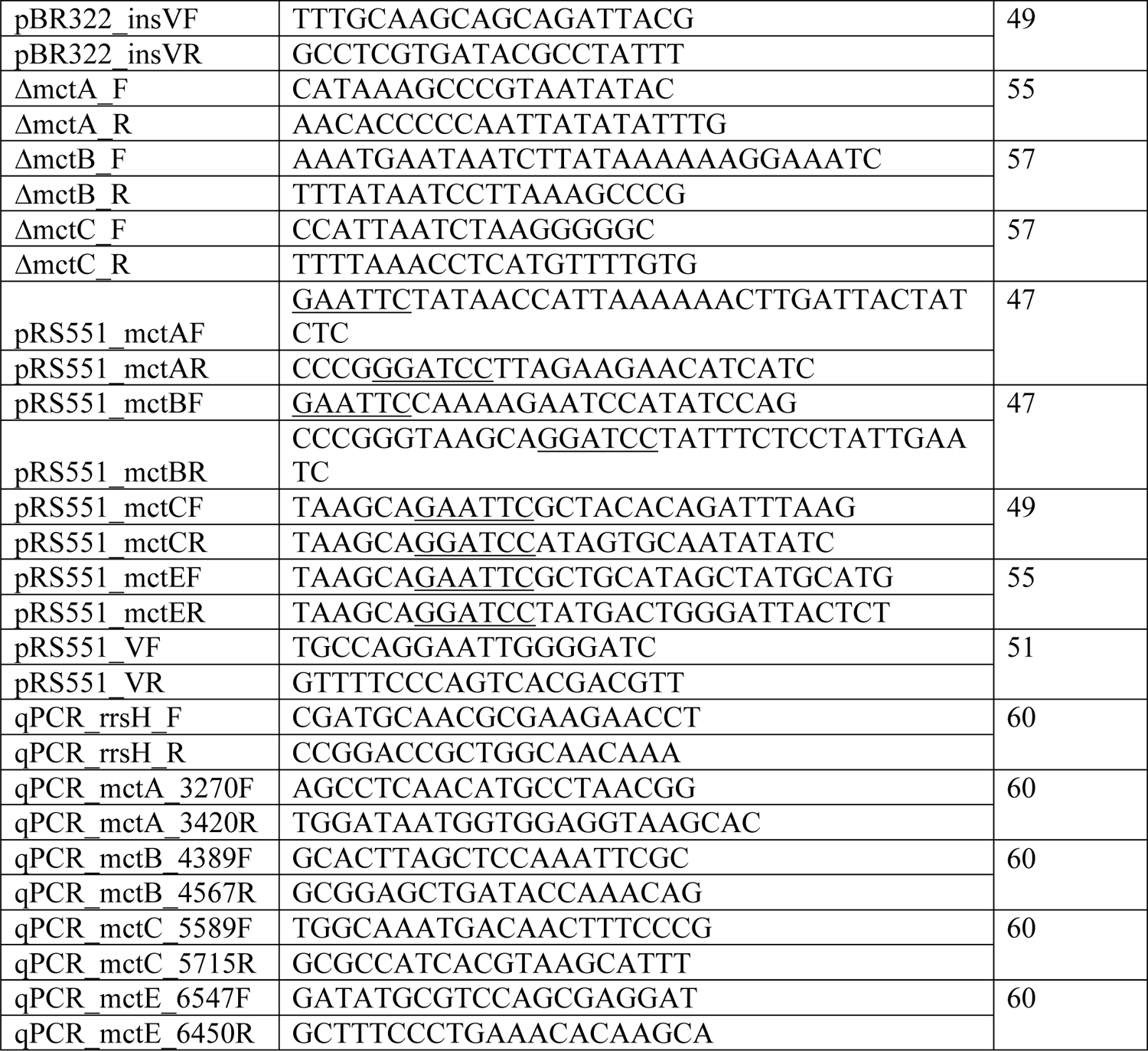
Strains, primers, and plasmids.

### Mcc1229 entry requires TonB

CirA is a known TonB-dependent transporter, and the import of CirA-dependent colicins requires the activity of TonB in the periplasm (45). Earlier data also implicated TonB in SOS induction by Mcc1229 in a reporter strain (15). We next sought to inactivate *tonB* in *E. coli* O157:H7 to determine its role in Stx2a amplification by Mcc1229. Attempts to delete *tonB* in the PA2 background by various methods were unsuccessful. *tonB* was instead deleted by one-step recombination in EDL933, a well-characterized *stx_1a_*^+^*stx_2a_*^+^ O157:H7 strain (46). A Δ*tonB*::*cat* mutant did not amplify Stx2a in response to supernatants containing Mcc1229 (Figure 6). When complemented with a plasmid copy of *tonB* (pKP315), the strain behaved as wildtype (Figure 6). These data indicated that CirA and TonB were both necessary for Stx2a amplification by Mcc1229.

**Figure 6.**
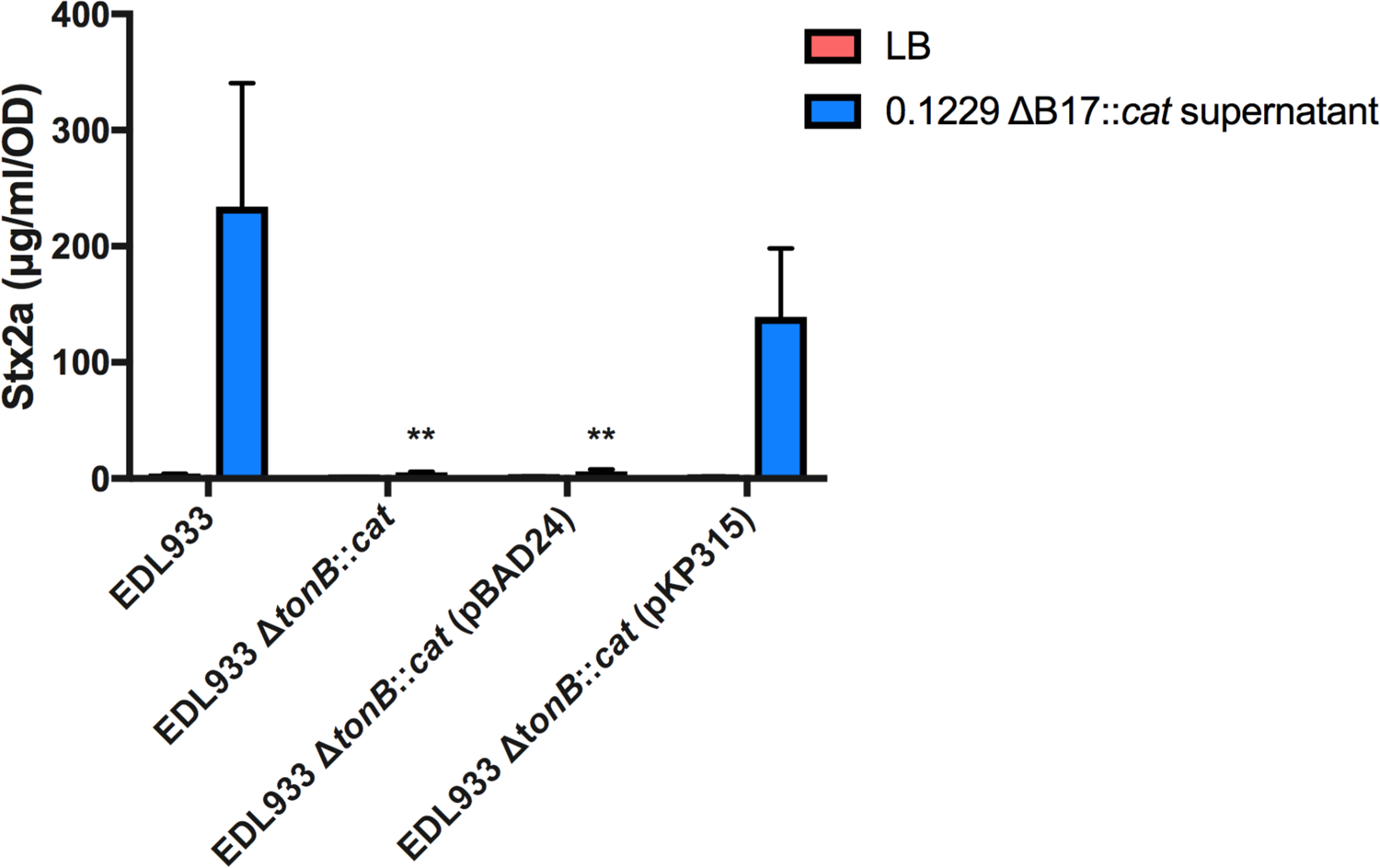
TonB is required for Mcc1229 activity. The periplasmic energy transducing protein TonB was deleted from *E. coli* O157:H7 strain EDL933 by one-step recombination. The resulting mutant does not increase Stx expression in response to spent supernatants containing Mcc1229. Activity is restored upon complementation with a plasmid copy of *tonB*, carried on pKP315. Because *tonB* is under P_araC_ control on pKP315, all strains in this experiment were grown in the presence of arabinose. Asterisks mark significant difference between a given supernatant sample and the EDL933 wildtype (two-way ANOVA, Dunnett’s multiple comparisons test).

### Immunity to Mcc1229 is mediated by the mctD-mctE region of p0.1229_3

The lethality of colicins and microcins necessitates a mechanism of protection for the producing cell. By cloning progressively smaller fragments of p0.1229_3 into pBR322, we identified a region of the plasmid that was sufficient to confer immunity to Mcc1229. Two adjacent ORFs, *mctD* and *mctE*, each predicted to encode a domain of unknown function (DUF)-containing protein, protected MG1655 from Mcc1229-mediated killing (Figure 7a). Vectors containing either of the single ORFs were not protective (Figure 7a). When the pBR322::*mctDE* construct was transformed into PA2, PA2 became insensitive to 0.1229 ΔB17::*cat* supernatant and Stx2a production did not increase (Figure 7b).

**Figure 7.**
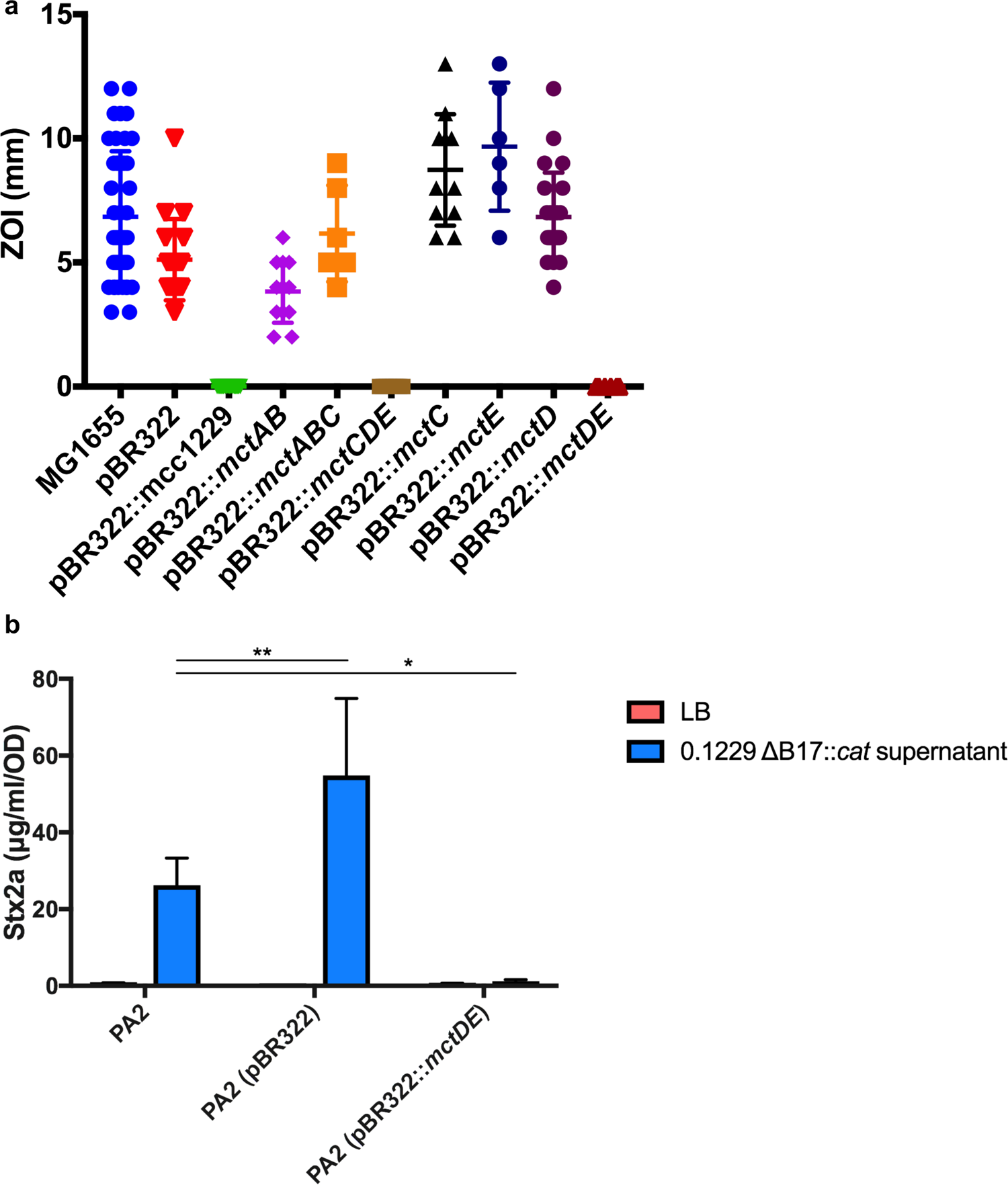
The *mctD*-*mctE* region is sufficient to confer immunity to Mcc1229. (a) Two p0.1229_3 ORFs annotated as proteins with domains of unknown function were cloned into pBR322 and transformed into *E. coli* strain MG1655. MG1655 carrying an empty vector is sensitive to Mcc1229 produced by *E. coli* 0.1229 Δ*B17*::*cat*. MG1655 carrying pBR322::*mctDE* is fully resistant to the microcin. (b) Transformants of PA2 carrying pBR322::*mctDE* do not increase Stx expression when grown in spent supernatants of *E. coli* 0.1229 Δ*B17*::*cat*. Asterisks mark significant difference between a given supernatant sample and the PA2 wildtype (two-way ANOVA, Dunnett’s multiple comparisons test; *, p < 0.05; **, p < 0.01).

### Mcc1229 is expressed in vivo but is not required for suppression of PA2

To determine the effect of microcins *in vivo*, we colonized germfree mice with *E. coli* 0.1229 and its derivatives and collected fecal samples at one day post infection. After suspending feces in LB, samples were centrifuged to pellet the solid matter and the supernatant spotted atop a suspension of the PA2 test strain. Supernatants from mice infected with 0.1229 inhibited the growth of PA2, but those from mice infected with a Mcc1229 knockout strain had no effect (Figure 8a and 8b).

**Figure 8.**
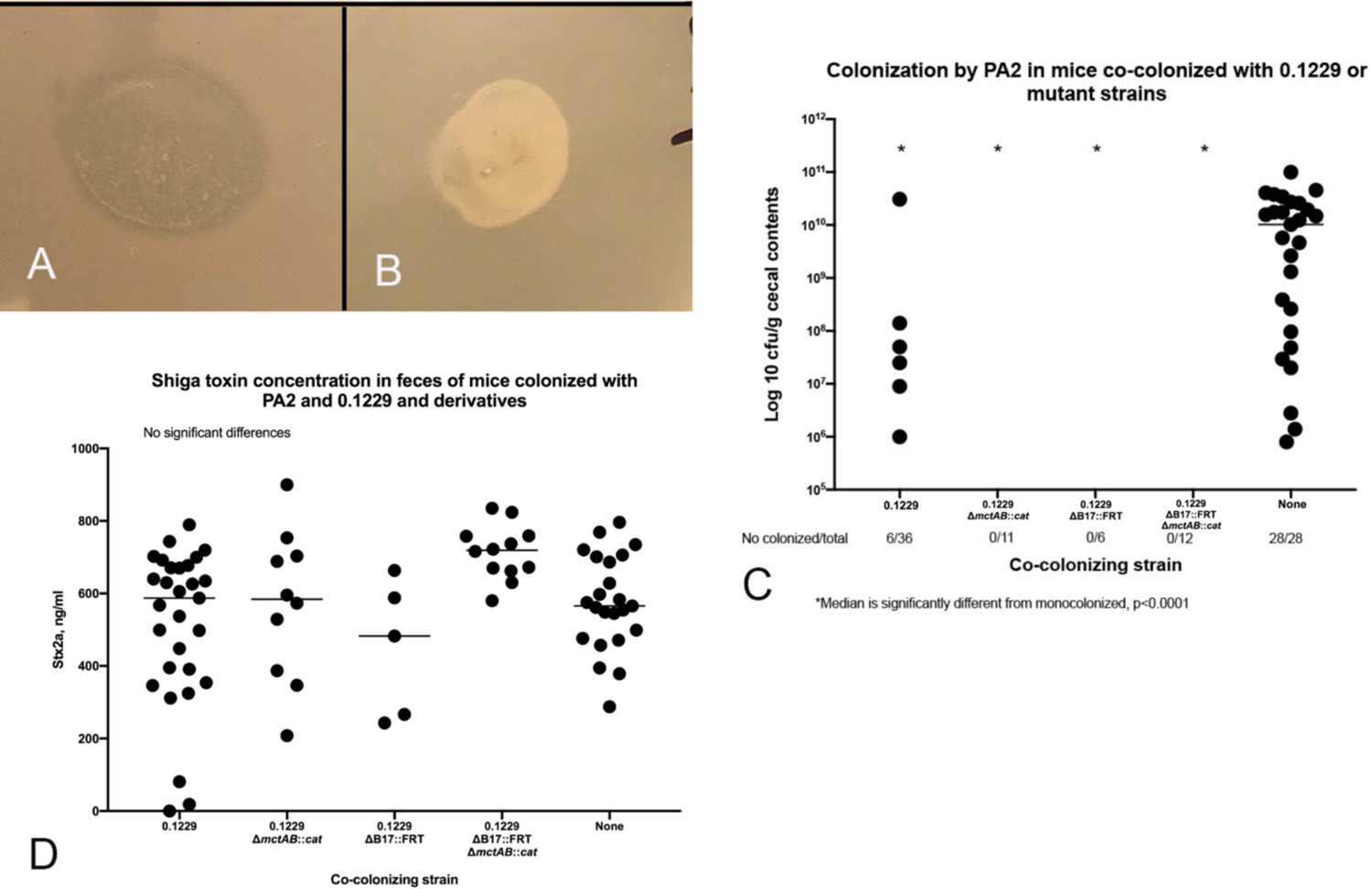
(a) and (b) Soft agar containing a suspension of PA2 was overlaid on LB, allowed to solidify, and fecal supernatants from infected mice were spotted on the overlay. Plates were incubated overnight at 37°C. (a) Fecal supernatants from mice co-colonized with 0.1229 and PA2 prevented growth of PA2, resulting in a zone of clearing in the soft agar. (b) Supernatants from mice colonized with 0.1229 ΔB17::FRT Δ*mctAB*::*cat* did not affect PA2 growth. (c) PA2 colonization in co-colonized mice. PA2 colonization was detectable in only 16% of mice co-colonized by wild-type 0.1229 and in none of the mice colonized by 0.1229 Δ*mctAB*::*cat*, 0.1229 ΔB17::FRT, or 0.1229 ΔB17::FRT Δ*mctAB*::*cat.* In contrast 100% of mice inoculated with PA2 alone became colonized (also see Table 2). (d) Low colonization level did not affect Stx production. Stx was detected in all mouse groups and concentration ranged from 18 to 900 ng/ml. There were no differences between groups.

**Table 2.**
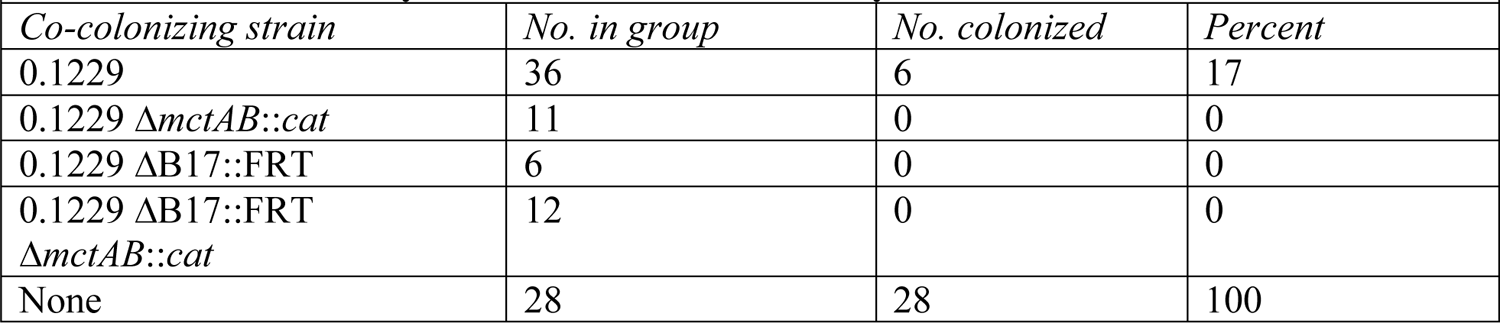
Colonization by PA2 in mice co-colonized by 0.1229 and derivatives.

| *Co-colonizing strain* | *No. in group* | *No. colonized* | *Percent* |
| --- | --- | --- | --- |
| 0.1229 | 36 | 6 | 17 |
| 0.1229 ΔmctAB::cat | 11 | 0 | 0 |
| 0.1229 ΔB17::FRT | 6 | 0 | 0 |
| 0.1229 ΔB17::FRT<br>ΔmctAB::cat | 12 | 0 | 0 |
| None | 28 | 28 | 100 |

The role of Mcc1229 in the germfree mouse model of EHEC was investigated by sequential inoculation of 0.1229 and PA2. Mice were first infected with 0.1229 or its derivatives, then with PA2 seven days later. Monoinfections of 0.1229 or PA2 served as controls. PA2 alone was able to colonize at concentrations of between 10^8^ – 10^10^ CFU/g and caused symptoms consistent with Stx-mediated disease including colitis and acute kidney injury (47, 48). When mice were colonized with 0.1229 prior to the introduction of PA2, PA2 colonization was almost fully suppressed. PA2 was recovered from the cecal contents of only six of 65 mice coinfected with 0.1229 or its derivatives (Figure 8c, Table 2). This effect did not require Mcc1229 or MccB17, however, as the single and double microcin mutants of 0.1229 were capable of suppressing PA2 equivalent to the wildtype (Figure 8c, Table 2). Colonization suppression was not protective against disease, as PA2 was still lethal to coinfected mice and Stx was detected in feces from all groups (Figure 8d, Table 3). This likely indicates that PA2 was present at some time during infection but was either lost or suppressed below the limit of detection.

**Table 3.**
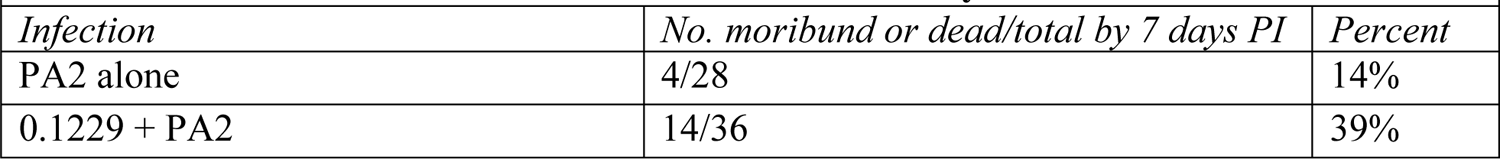
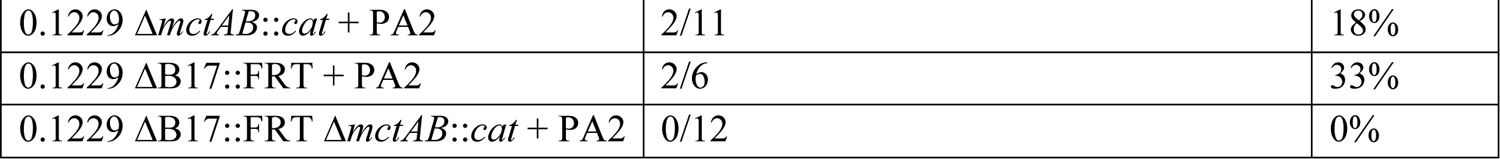
Clinical illness due to PA2 in mice co-colonized by 0.1229 or its derivatives.

| *Infection* | *No. moribund or dead/total by 7 days PI* | *Percent* |
| --- | --- | --- |
| PA2 alone | 4/28 | 14% |
| 0.1229 + PA2 | 14/36 | 39% |
| 0.1229 $\Delta mctAB::cat$ + PA2 | 2/11 | 18% |
| 0.1229 $\Delta B17::FRT$ + PA2 | 2/6 | 33% |
| 0.1229 $\Delta B17::FRT \Delta mctAB::cat$ + PA2 | 0/12 | 0% |

## Discussion

A putative microcin from the human *E. coli* isolate 0.1229 was previously shown to induce the SOS response and Stx expression in target strains (15). Here, we have confirmed the activity of this microcin (Mcc1229), isolated its activity from that of a second microcin encoded by 0.1229 (MccB17), and further characterized its production, regulation, and effects. Like several other colicins and microcins, Mcc1229 uses the CirA siderophore receptor (Figure 5) and the TonB complex (Figure 6) for entry into a target cell. CirA was first identified as the <u>c</u>olicin <u>I receptor</u> and is also used by colicin/microcin V (33, 49). In the producing strain, evidence suggests that Mcc1229 requires the *mctABC* region of plasmid p0.1229_3 for activity (Figure 3b). Open reading frames similar to *mctA*, with cysteine-rich C-terminal regions and cognate ABC transporters, are also consistent with typical microcin operons (50).

The functional contributions of the cupin-like *mctC* and DUF-containing *mctD*-*mctE* ORFs in the Mcc1229 cluster have not yet been elucidated. In our system, the *mctDE* region conferred immunity to Mcc1229 killing and Stx amplification (Figure 7), but it is not clear whether the DUF-containing proteins encoded by *mctD* and *mctE* directly interact with the microcin. Current Pfam records indicate that the DUF2164 domain present in MctE is found in 804 protein sequences in 715 species, but it is not associated with a clan or superfamily (51). DUF4440, which is found in MctD, belongs to a family in the nuclear transport factor (NTF) 2 clan, which includes numerous proteins with enzymatic and non-enzymatic functions (52). Some proteins with NTF2-like folds are known to provide immunity to bacterial toxins, but their sequences (Pfam PF15655) are diverse and dissimilar to the DUF4440 domain in MctD (53). Proteins with DUF4440 and/or NTF2-like domains have also been shown to operate in polyketide biosynthesis pathways, where they are involved in catalyzing the formation of natural products (54, 55). Some proteins with cupin domains have enzymatic activity (56, 57), so it is possible that the p0.1229_3 MctC is involved in processing or modification of Mcc1229. Because our attempts to complement the *mctA*, *mctB*, and *mctC* deletion mutants *in trans* were unsuccessful (data not shown), we cannot speculate further on the contributions of these ORFs to Mcc1229 production.

Beyond its cellular export and import, the observed Fe-Fur regulation of Mcc1229 further supports its classification as a microcin. Mcc1229’s amplification of Stx was increased in the presence of chelating agents, and this effect could be reversed by the addition of iron specifically (Figure 2). Moreover, the expression of *mctA*, *mctB*, *mctC,* and *mctE* genes were increased in a Δ*fur*::*cat* background (Figure 4b). Taken together, these data likely indicate that Mcc1229 is transcriptionally repressed by the canonical Fe-Fur complex (58). A similar pattern is seen in the regulation of microcin E492 in *Klebsiella pneumoniae* (29). Like the site upstream of *mceX* in the MccE492 operon, the putative Fur box upstream of *mctA* is 68% (13/19 nt) identical to the consensus Fur sequence described for *E. coli* (59, 60). Fur-regulated microcins may provide a competitive advantage for *E. coli* strains *in vivo*, where iron availability is restricted (61, 62). The Fur regulon is essential for survival of various *E. coli* pathotypes *in vivo* (63–65), and the B2 phylogroup of *E. coli* (which includes 0.1229) is associated with a high prevalence of microcin genes (66).

In our study, microcin Mcc1229 was produced *in vivo* but had no effect on EHEC colonization or disease (Figure 8). Nevertheless, we observed a striking example of suppression by *E. coli* 0.1229 in which PA2 was rarely if ever recovered from coinfections. Most other *E. coli* do not suppress EHEC to the same extent, although there is precedent for colonization suppression by the probiotic strain Nissle 1917 (67, 68). Intriguingly, Nissle and 0.1229 both belong to ST73, a lineage frequently isolated from ExPEC infections (69). ST73 strains carry a broad assortment of virulence factors, including many genes for adherence and iron acquisition that could provide a selective advantage over competitors (70). Ongoing studies may determine whether colonization resistance is a trait that is common to ST73.

Interactions with the microbiome can alter the virulence of EHEC in numerous ways. Understanding these effects will help predict the unique pathogenicity and disease outcomes of a given infection. Here, we have expanded upon the attributes of Mcc1229, a new *E. coli* microcin that induces the SOS response and amplifies Stx2a expression *in vitro*. When characterizing the interplay of Mcc1229 and EHEC *in vivo*, however, we found that microcin activity was not a significant contributor to EHEC virulence or to colonization efficiency. This discrepancy highlights the need for additional research regarding the dynamics of bacteriocin expression in the intestinal environment. The regulation, stability, and activity spectrum of bacteriocins all influence their physiological role, as do external factors such as inflammation and nutrient availability. Although Mcc1229 could be unified with other microcins based on the cellular factors described in this work, its actual ecological impact was not apparent from our germfree mouse model and awaits further clarification.

## Materials and Methods

### Bacterial strains and culture conditions

*E. coli* strains were routinely grown in lysogeny broth (LB; 10 g/l tryptone, 5 g/l yeast extract, 10 g/l NaCl) at 37°C and maintained in 20% glycerol at −80°C. Minimal medium (M9) was formulated with 12.8 g/l Na_2_HPO_4_*×*7H_2_O, 3 g/l KH_2_PO_4_, 0.5 g/l NaCl, 1 g/l NH_4_Cl, 2 mM MgSO_4_, and 0.1 mM CaCl_2_. M9 was supplemented with 0.1% casamino acids, 0.005% thiamine, and 0.4% of the desired carbon source. Mueller-Hinton (MH) agar was prepared according to the manufacturer’s instructions. EDTA, 2,2’-bipyridyl, FeCl_3_, CaCl_2_, MgCl_2_, and MnCl_2_ were added to media at 0.2 mM. Antibiotics were used as follows: ampicillin, 50 µg/ml; chloramphenicol, 12.5 µg/ml; kanamycin, 25 µg/ml; tetracycline, 10 µg/ml. All media components were purchased from BD Difco (Franklin Lakes, NJ) and all enzymes from New England Biolabs (NEB; Ipswich, MA) unless otherwise noted.

### One-step recombination

*E. coli* knockouts were constructed according to the protocol of Datsenko and Wanner (71). Primers incorporating 40 bp immediately upstream and downstream of the gene of interest were used to amplify the *cat* cassette from pKD3 or the *kan* cassette from pKD4 (Table 1). The target strain was first transformed with pKD46 and grown to mid-log phase, then induced with 0.02 M L-arabinose for 1 h. Cells were washed with cold water and 10% glycerol and electroporated with the *cat* or *kan* PCR product using a GenePulser II instrument (2.5 kV, 0.2 cm gap cuvettes; Bio-Rad, Hercules, CA). Transformants were verified by colony PCR with primers approximately 200 bp up- and downstream of the gene of interest, and the site of the insertion was confirmed by Sanger sequencing (Table 1). Mutants were complemented with a plasmid copy of the gene of interest, cloned into the medium-copy number vector pBR322 by Gibson assembly (72). Assembly primers were designed using NEBuilder (https://nebuilder.neb.com; Table 1). Amplicons were purified with the QIAquick Cleanup Kit (Qiagen, Germantown, MD) and assembled with the Gibson Assembly Cloning Kit according to the manufacturers’ instructions. Assembly junctions were likewise confirmed with colony PCR and Sanger sequencing (Table 1).

Because multiple efforts to inactivate *tonB* in PA2 by one-step recombination were unsuccessful, we generated a Δ*tonB*::*cat* mutant in the EDL933 background. This mutant was complemented by pKP315, kindly provided by Dr. Kathleen Postle, which carries an arabinose-inducible copy of *tonB*. L-arabinose was added to EDL933 cultures at 0.3%.

### LacZ fusions

Transcriptional activity was measured by fusing selected p0.1229_3 fragments to a promoterless *lacZ* gene in the pRS551 vector (73). Fragments were amplified from p0.1229_3 using the given primers (Table 1) and digested with EcoRI-HF and BamHI-HF enzymes. The products were cleaned up using the QIAquick kit and ligated into an EcoRI-BamHI digest of pRS551. Ligation mixtures were transformed into chemically competent DH5*α* cells (New England Biolabs) and verified by miniprep and restriction digests. Constructs were then electroporated into *E. coli* 0.1229. Reporter strains were cultured in LB, shaking at 37°C, and grown until mid-logarithmic phase. Cells were then harvested and suspended in Z buffer. LacZ activity was measured by the hydrolysis of *o*-nitrophenyl-β-D-galactoside according to the method of Miller *et al.* (74).

### qPCR

RNA was extracted from 16 h LB cultures of 0.1229 and 0.1229 Δ*fur*::*cat* using TRIzol (ThermoFisher Scientific, Waltham, MA). Genomic DNA was removed by digestion with RQ1 RNase-Free DNase (Promega, Madison, WI) and RNA converted to cDNA using the ThermoScript RT-PCR system (ThermoFisher). Expression of *mctA*, *mctB*, *mctC*, and *mctE* genes was quantified in 20 µl reactions using PerfeCTa SYBR Green FastMix (Quantabio, Beverly, MA) and 200 nM qPCR primers (Table 1) on a QuantStudio3 instrument (ThermoFisher). To validate the efficiency (>95%) of each primer pair, its target was amplified from genomic DNA and purified in a spin column cleanup kit (Dot Scientific Inc., Burton, MI). The concentration of this product was measured by spectrophotometry (NanoDrop 1000, ThermoFisher) and ten-fold dilutions ranging from 10^-2^ through 10^-7^ ng/µl were used as templates in qPCR. A standard curve was constructed from the resulting Ct values. Differences in gene expression between wildtype and Δ*fur*::*cat* strains were determined by the ΔΔCt method, using the 16S ribosomal RNA *rrsH* gene as an internal control (75).

### In-frame deletions and site-directed mutagenesis

Prior work demonstrated that a fragment of the p0.1229_3 plasmid encompassing nucleotides 2850 through 7950 was sufficient to amplify Stx when cloned into pBR322 (15). In-frame deletions of individual ORFs in this vector, pBR322::mcc1229, were generated with NEB’s Q5 Site-Directed Mutagenesis Kit. Primers facing outward from the chosen ORF were designed with the NEBaseChanger tool and used with Q5 polymerase to amplify a linear fragment from pBR322::mcc1229 (Table 1). This product was treated with KLD enzyme cocktail to digest template DNA and recircularize the plasmid according to the manufacturer’s instructions. Constructs were confirmed by PCR of DH5*α*transformant colonies using VF/VR primers (Table 1). Mutations were then verified by Sanger sequencing and plasmids electroporated into C600 as described above to assure that no wildtype copies remained.

### Inhibition assays

Microcin production was evaluated by measuring inhibition of a target strain in agar overlays (76). The microcin-producing strain was spot-inoculated on MH agar and incubated at 37°C for approximately 24 h. Plates were inverted over filter paper discs impregnated with 300 µl chloroform for 30 minutes to kill producing cells. Cultures of the target strains were then suspended to 0.05 OD_600_ per ml in soft (0.7%) nutrient agar, poured atop the plates, and allowed to solidify. After overnight incubation at 37°C, inhibition was noted by the presence of halos surrounding a microcin-producing colony. Zones of inhibition were quantified by subtracting the diameter of the producing colony from the diameter of the clear zone surrounding it. Spontaneous mutants growing within the zones of inhibition were restreaked to purify and retested in agar overlays to confirm microcin resistance. Known microcin and colicin producers and their corresponding indicator strains were from the NCTC reference set, kindly provided by Dr. Robert *F.* Roberts (44).

For inhibition assays using supernatants, plates were inoculated with the test strain in soft agar as described above. Fecal samples from mice colonized with 0.1229 or its derivatives were collected at 1 day after inoculation with PA2, suspended in 100-200 µL LB broth, and centrifuged. Ten µl of supernatant were spotted atop the test strain and allowed to dry before overnight incubation at 37°C.

### Whole-genome sequencing and bioinformatics

Genomic DNA was extracted from overnight cultures using the DNeasy Blood and Tissue Kit (Qiagen). Libraries were prepared using the Nextera XT kit (Illumina, San Diego, CA) and sequenced on the MiSeq platform, generating 2*×*250 bp reads. Reads were assembled in the Galaxy workspace with the SPAdes tool (77), and single nucleotide polymorphisms were identified using Snippy (78), comparing to the reference genome assembly GCA_000335355.2. Putative Fur binding sites and promoter motifs were identified by analysis of the p0.1229_3 sequence with RSAT (79) and BPROM (80), respectively.

### Coculture and supernatant experiments

Supernatants of *E. coli* 0.1229 and its derivatives were harvested after 16 h shaking at 37°C and passed through 0.2 µm cellulose acetate filters (VWR Life Sciences, Radnor, PA). Assays to quantify Stx amplification were performed as previously described (15). Briefly, the test strain of *E. coli* was suspended in 1 ml of spent supernatant to 0.05 OD_600_ and inoculated atop solid LB agar in a 6-well plate (BD Biosciences Inc., Franklin Lakes, NJ).

For Stx assays, the test strains were *E. coli* O157:H7 isolates. PA2 (81) was used routinely as it demonstrated the greatest Stx amplification in prior experiments (13). EDL933 (46) was used in the event that a PA2 mutant could not be obtained. Strains were diluted to 0.05 OD_600_ in either broth or filtered supernatant and inoculated atop solid LB agar in 6-well plates. Cultures were then incubated statically at 37°C for 8 h. Aliquots of each culture were removed to measure OD_620_, and the remaining volume was treated with 6 mg/ml polymyxin B for 5 min at 37°C to release intracellular Stx. Samples were then centrifuged for 5 min to pellet cell debris and supernatants were collected and stored at −80°C until use in an R-ELISA.

### R-ELISA

Shiga toxin was detected in a receptor-based ELISA as previously described (14). A microtiter plate was first coated with 25 µg/ml ceramide trihexosides (Matreya Biosciences, Pleasant Gap, PA) in methanol. The methanol was evaporated and the plate subsequently blocked with 4% bovine serum albumin in phosphate buffered saline containing 0.05% Tween 20 (PBST). Supernatant samples were diluted in PBS as necessary and added to wells for 1 h, gently shaking at room temperature. Monoclonal anti-Stx2 antibody was purchased from Santa Cruz Biotech (Santa Cruz, CA) and added to wells at 1 µg/ml for 1 h. Anti-mouse secondary antibody conjugated to horseradish peroxidase was purchased from MilliporeSigma (Burlington, MA) and also added at 1 µg/ml for 1 h. Between each of the preceding steps, the plate was washed five times with PBST. One-step Ultra TMB (ThermoFisher) was then used for detection. The plate was incubated for approximately 5 min before the reaction was stopped with the addition of 2 M H_2_SO_4_ and the A_450_ was measured (Multiskan FC, ThermoFisher). A standard curve was established using serial dilutions of lysate from PA11, a high Stx2a-producer (81). The concentration of Stx2a in *E. coli* O157:H7 samples was determined by comparison to this curve and is reported in µg/ml, normalized to the OD_620_ of each *E. coli* O157:H7 culture.

### Animal experiments

Male and female Swiss-Webster mice aged 3 to 5 weeks were raised in the University of Michigan germfree colony. They were housed in soft-sided bubble isolators or sterile isocages and fed autoclaved water and laboratory chow *ad libitum*. Throughout the experiment, the mice received sterile food, water, and bedding to maintain germfree conditions, except for the infecting *E. coli* strains. All animal experiments were conducted with the approval of the University of Michigan Animal Care and Use Committee.

Mice were infected orally with ∼10^6^ CFU of each *E. coli* inoculum. In coinfection experiments, 0.1229 and its derivatives were inoculated first and were followed by PA2 one week later. Mice were weighed prior to each inoculation and just prior to euthanasia. They were evaluated daily for evidence of illness (dehydration, ruffled coat, or reluctance to move) and were euthanized 1 or 7 days after PA2 infection or when they became moribund. Prior to euthanasia, evidence of illness was recorded, and at necropsy, samples were collected for bacterial culture, Stx2 ELISA, and histologic examination.

For bacterial culture, samples of the cecal contents were weighed, serially diluted in sterile LB broth, and cultured on sorbitol-MacConkey (SMaC) agar. PA2 is non-sorbitol-fermenting and appears as white colonies on SMaC plates. Cultures from co-colonized mice were quantified based on the number of pink or white colonies. For quantification of Stx2, the cecal contents were stored at −20°C until evaluation with a Premier EHEC ELISA kit (Meridian Biosciences Inc., Cincinnati, OH). The concentration of Stx2a was determined by comparison to the PA11 standard curve discussed above (81).

## Acknowledgments

This work was supported by grant 1 R21 AI130856-01A1 to E.G.D. by the National Institute of Allergy and Infectious Diseases and by USDA National Institute of Food and Agriculture Federal Appropriations to E.G.D. under project PEN04464 and accession number 1015714.

## Author Statements

The authors declare that there are no conflicts of interest.

